# Co-option of a seed-like proteome by oil-rich tubers

**DOI:** 10.1101/2020.12.15.422834

**Authors:** Philipp William Niemeyer, Kerstin Schmitt, Oliver Valerius, Gerhard H. Braus, Jan deVries, Anders Sven Carlsson, Per Hofvander, Till Ischebeck

## Abstract

Co-option is an important aspect of evolution that can occur on several levels. Genes, whose function was molded by selection in the evolutionary past, are readily observed to serve a new function when acting in a different context in an extant system. Whole organs can be co-opted for new roles as well. For example, roots that evolved from shoot-like axes. Finally a framework of genes and its coded proteins can be co-opted to serve a similar molecular function but in a completely different organ, drastically changing its properties. Here, we describe such an example, where a set of proteins important for desiccation tolerance and oil accumulation in seeds of most angiosperms was co-opted in the tubers of yellow nutsedge (*Cyperus esculentus*). These tubers are not only desiccation tolerant but also store a large amount of lipids—especially TAG, similar to seeds. We generated nanoLC-MS/MS-based proteomes in five replicates of four stages of tuber development and compared them to the proteomes of roots and leaves, yielding 2257 distinct protein groups. Our data reveal a striking upregulation of hallmark proteins of seeds in the tubers. A deeper comparison to a previously published proteome of Arabidopsis seeds and seedlings indicate that indeed a seed-like proteome was co-opted. This was further supported by an analysis of the proteome of a lipid-droplet enriched fraction of yellow nutsedge, which also displayed seed-like characteristics.

## Introduction

Drought is the prime stressors that land plants have to overcome since their dawn (1). One strategy that land plants employ is to limit water loss and to transport water via a vascular system (2, 3). Another is to produce desiccation tolerant cells and tissues (4). Common to drought tolerance and desiccation is the accumulation of small osmolytes and proteins with protective functions (4). Also common to both strategies is the accumulation of neutral lipids, foremost triacylglycerol (TAG), in cytosolic lipid droplets (LDs) with especially high levels being reached in embryonic tissues (5). In most flowering plants, desiccation tolerance is limited to seeds and to some extend pollen. However, there are also plants that have desiccation tolerant vegetative organs some of the also referred to as “resurrection plants” such as *Sporobolus stapfianus* (6) and *Craterostigma plantagineum* (7) and it is likely that this tolerance evolved several times independently (8). It is however likely that these independent origins of desiccation tolerance are underpinned by the co-option of existing regulatory programs for resilience (9).

Here, we investigated yellow nutsedge (*Cyperus esculentus*), a monocot, perennial C4 plant (10). This species produces stolon-derived underground tubers that can fully desiccate and remain viable for years (11). Furthermore they can accumulate 25-30 % of their dry mass in lipids especially TAG (11, 12). In this regard they are unique as even the tubers of a close relative, purple nutsedge (*Cyperus rotundus*), are neither desiccation resistant nor oil accumulating (13, 14). We studied the proteomes of yellow nutsedge tubers and compared them to previously published proteomes of Arabidopsis seeds and seedlings. This comparison revealed striking similarities between tubers and seeds, and tuber sprouting and seedling establishment. We conclude that this species evolved the desiccation tolerance and oil-richness in their tubers by co-opting a protein framework normally present predominantly in seeds.

## Results and Discussion

### Tubers of yellow nutsedge are enriched in hallmark proteins of seeds

To understand the molecular basis of tuber resilience in yellow nutsedge, we studied the proteome of four stages of tuber development (freshly harvested, dried, rehydrated for 48 h, and sprouted), and compared it to roots and leaves (Figure S1). From the first three tuber stages, we additionally isolated LD-enriched fractions. After a tryptic digest, all peptide samples were analyzed by LC-MS/MS in five biological replicates and all protein groups were quantified using a label-free MS1-based quantification algorithm resulting in relative iBAQ (riBAQ) intensities. For functional annotation of these proteins, we used all nutsedge library entries as query for a BLASTp against the Arabidopsis TAIR10 primary transcript protein release library (15).

First, we studied the total proteomes. A principal component analysis (PCA) revealed that the tuber proteomes were mostly separated by developmental stage but much more distinct from roots and leaves (Figure 1A). In order to identify tuber-specific proteins, the data was hierarchically clustered (Figure 1A). Nine of the resulting thirty clusters (clusters 3-6, 8-11 and 13) contained 477 protein groups highly enriched in at least one of the tuber stages and far less abundant in leaves and roots. These protein groups corresponded to 433 Arabidopsis homologs that were subjected to a gene ontology search (16). This search indicated that 54 of these enriched proteins were associated with carbohydrate metabolism and 41 with cellular responses to stress but similar numbers were found in the other clusters combined. Overrepresented, however, were starch metabolism and several seed-related processes including seed maturation, dormancy, germination and seedling development. None of these seed-related terms were overrepresented in the other combined clusters.

**Figure 1.**
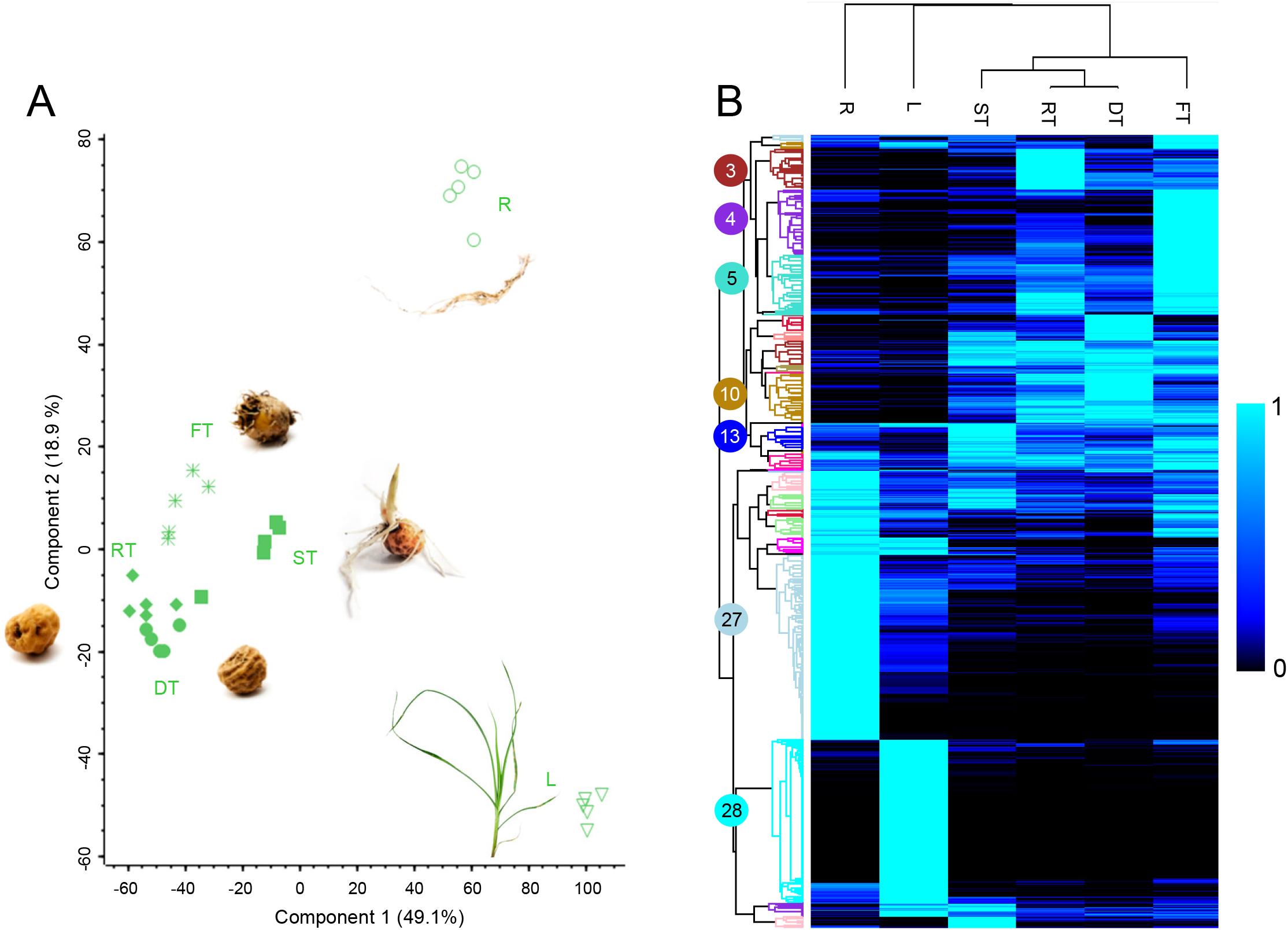
Developmental tuber stages have a distinct proteome. Comparative proteomics of total protein extracts of tubers (fresh, FT; dry, DT, rehydrated for 48 h (RT) and sprouted (ST), leaves (L) and roots (R; Table S2). (A) Principal component analysis plot. (B) The values were normalized setting the average of the stage with the highest abundance to 1 and these values were hierarchically clustered. Proteins sorted by cluster can be found in (Table S3A). n=5 biological replicates per stage.

Seven of the ten most abundant proteins that were at least tenfold enriched in one of the tuber stages in comparison to leaves and roots (Tables 1) have Arabidopsis homologs that are almost exclusively expressed in seeds based on transcript data (17); these homologs are further strongly enriched in proteomes of Arabidopsis seeds compared to 60 h old seedlings (18). Data gleaned from the total proteome thus suggest that yellow nutsedge might co-opt a seed-like proteome to sustain desiccation tolerant oil rich tubers.

**Table 1.**
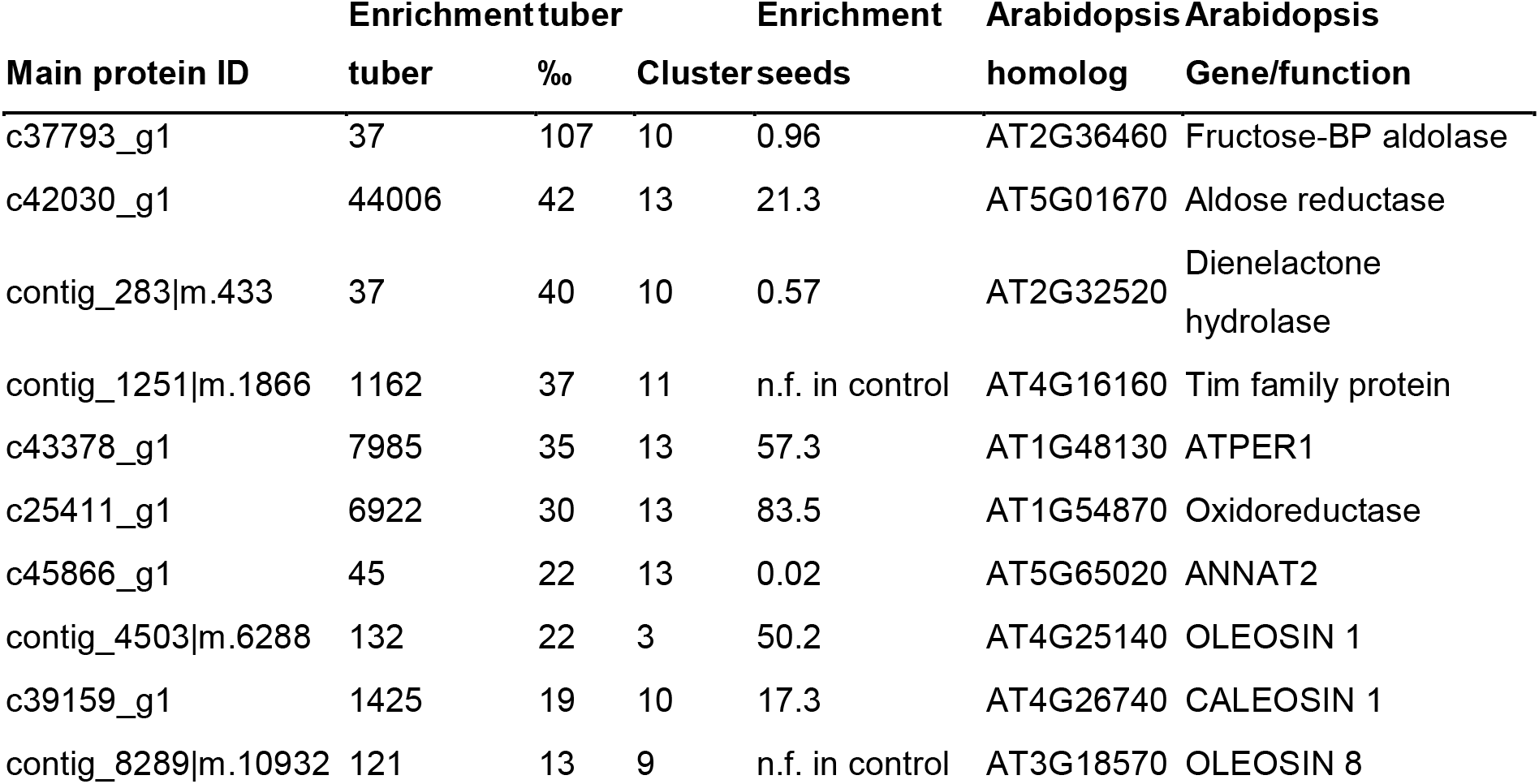
The majority of the most abundant proteins strongly enriched in tubers have seed-specific homologs in Arabidopsis. The protein compositions of yellow nutsedge tubers (fresh, dry, rehydrated and sprouted) were compared to the ones of leaves and roots. Displayed are the ten most abundant proteins found in the tuber samples that were at least tenfold enriched in comparison to leaves and roots (by highest average). Enrichment in seeds was calculated between rehydrated seeds and 60 h old seedlings (18). Arabidopsis homologs were determined by BLASTp analysis.

### Proteomes of tubers of yellow nutsedge and Arabidopsis seeds correlate

To further explore the similarity of tubers of yellow nutsedge and seeds of Arabidopsis, we compared the developmental pattern between the six yellow nutsedge stages/tissues investigated in this study with six stages of seedling establishment from Arabidopsis (18). 1200 protein homologs were present in the proteomes of both species and in all replicates of at least one stage. To allow for meaningful between-species comparisons, we defined, in each species and for each protein the highest average abundance in a given stage as 1. This dataset was then analyzed in three ways.

First, a PCA revealed that the tuber and seed samples clustered closely together with the exception of fresh tubers that were more closely related to seedlings (Figure 2A).

**Figure 2.**
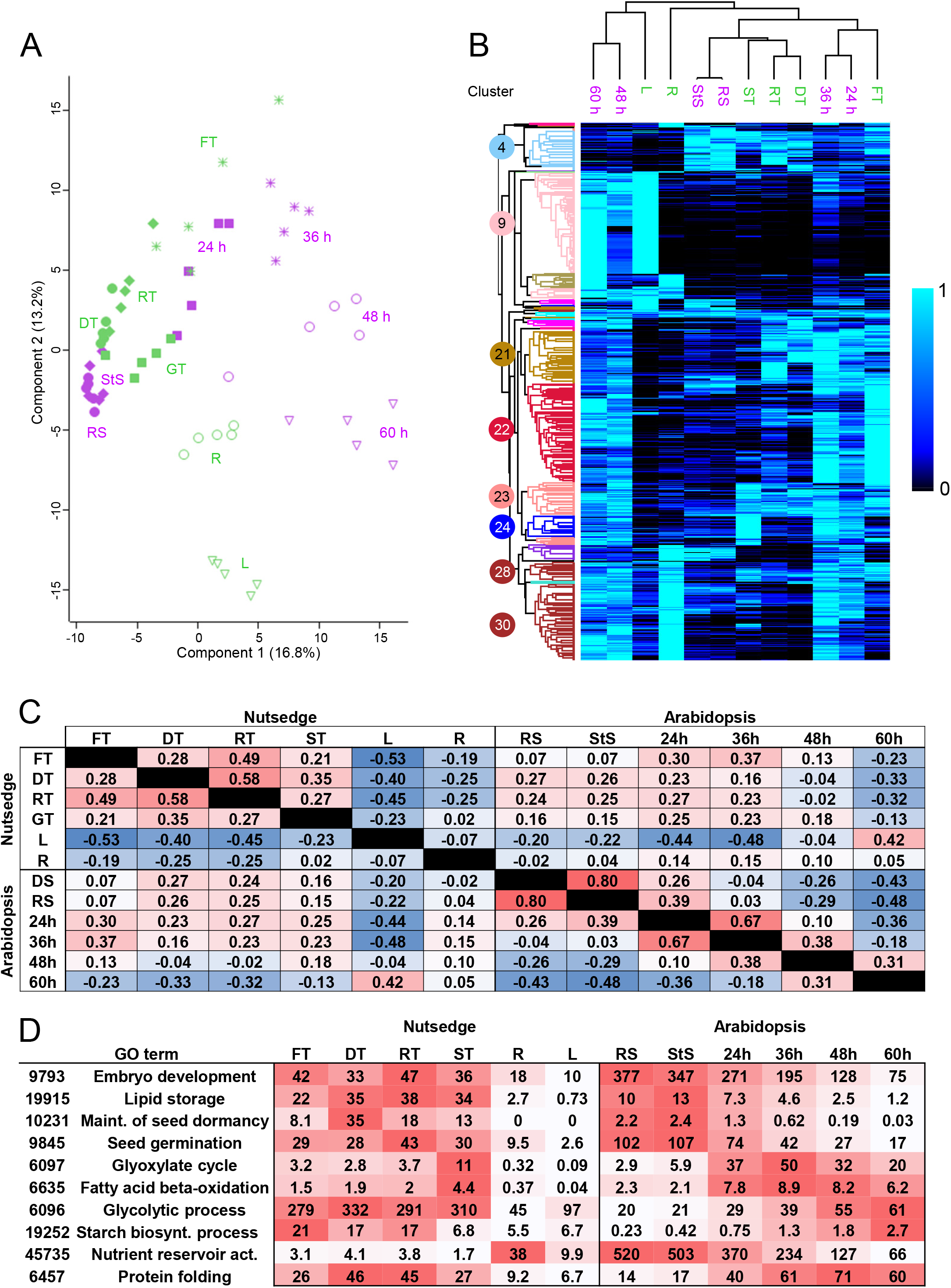
Comparison of protein expression pattern between nutsedge tissues and Arabidopsis seeds and seedlings. Total protein extracts of yellow nutsedge tubers (fresh, FT; dry, DT, rehydrated for 48 h, RT; and sprouted, ST), leaves (L) and roots (R) were compared to Arabidopsis seeds (rehydrated, RS, and stratified, StS) and seedlings (24 h – 60 h in the light after stratification). The principal component analysis (A) compares the individual samples. For the hierarchical clustering (B, see Table S5 for clusters) and the Pearson correlation analysis (C), samples were averaged for each stage. (D) Several GO terms are displayed that show similarities but also differences between tubers and seeds. Proteins were grouped by GO-terms and their relative abundance added up indicating per mill of the total proteome. A list of all GO terms can be found in Table S6 and S7. n=5 for all stages. Arabidopsis protein data was previously published in Kretzschmar et al. 2020.

Second, hierarchical clustering of the data (Figure 2B) yielded two clusters (4+5), comprising 90 proteins with a marked elevation in seeds rehydrated for 30 min and seeds stratified for 74 h at 4°C, and in dry and rehydrated tubers. These proteins included known LD-associated proteins such as oleosins and caleosins, late embryogenesis abundant (LEA) proteins and class I small heat shock proteins. All these protein families are known to be highly abundant in seeds (19-21) but likewise important in stress responses (22-25). Cluster 24 noteworthily contains 48 proteins high in sprouting tubers and seedlings, speaking to similarities between seedling establishment and tuber sprouting. Included in this cluster are proteins of the β-oxidation and glyoxylate cycle, needed for the conversion of lipids into carbohydrates. Further, tubers and seeds again clustered together.

Third, we calculated correlation coefficients between the averages of the different developmental stages to estimate similarities (Figure 2C). Here it was evident that, while tubers were most similar to each other, they also showed a positive correlation with the Arabidopsis seed and young seedling stages. Reversely, Arabidopsis seeds had higher correlation coefficients to nutsedge tubers than to 36 h old Arabidopsis seedlings.

### Homologs to proteins with important functions in seeds and seedlings are enriched in tubers

GO term analyses based on the total abundance of proteins bolstered the similarities of tubers and seeds (Figure 2D). For instance, the terms embryo development, lipid storage, maintenance of seed dormancy and seed germination were especially high in seeds and tubers but reduced in older stages. The term maintenance of seed dormancy only comprises one protein, a homolog to the Arabidopsis protein 1-CYSTEINE PEROXIREDOXIN 1 (PER1) that is enhances primary seed dormancy by suppressing ABA catabolism and GA biosynthesis (26). Another protein, homolog to GEM-RELATED 5 (GER5, AT5G13200), displays a similar pattern. GER5 is an ABA responsive gene that also regulates germination in Arabidopsis seeds (27). As of this, it can be speculated that ABA and other hormones could play a role in oil accumulation, desiccation tolerance, and dormancy in tubers—mirroring seeds (28).

β-Oxidation and the glyoxylate cycle are key during Arabidopsis seedling establishment. This was further reflected in GOterms of sprouting tubers—although overall not reaching the same levels as Arabidopsis does, especially in case of the glyoxylate cycle. Yellow nutsedge thus seems to utilize its oil to synthesize carbohydrates, despite storing additionally large amounts of sugars and starch that are also degraded during sprouting (29).

The GO term analysis, however, also displayed differences between tubers and seeds. Tubers had higher levels of proteins associated with starch biosynthesis and glycolytic processes, likely reflecting that the tubers store starch and free sugars in higher amounts (30). Another difference is the amount of storage proteins present. While members of cupin storage protein families are enriched in tubers and decrease during sprouting, Arabidopsis seeds contain a far greater amount of storage proteins amounting to more than half of the total protein based on the proteomic data; this amount is far lower in tubers (< 0.5 %). Tubers on the other hand contain many proteins associated with protein folding, including various heat shock protein families that might protect proteins during desiccation. In contrast, these types of proteins are less abundant in Arabidopsis seeds and accumulate only during seedling establishment indicating that here other strategies are employed to preserve protein integrity.

### Lipid droplet proteins known from seeds abound in tubers of yellow nutsedge

Most plant cells store TAG predominantly in cytosolic lipid droplets (LDs)(31). LDs are presumably present in all cell types but can strongly differ in their protein composition (18, 32-35). The described similarities between tubers and seeds and the notion that both accumulate oil raises the question if the tuber proteins that are associated with LDs are also similar to seeds. Therefore, we analyzed LD-enriched fractions of three tuber stages.

Homologs to known LD proteins made up ~60-70% of the proteins in LD-enriched fractions—with only ~5% being found in total extracts. With one exception, all these homologs were significantly enriched in the LD fraction (Figure 3A) indicating that these homologs are also LD associated. The most abundant among these proteins were the oleosins (Figure 3B)—especially the isoforms with higher sequence similarity to the Arabidopsis main seed oleosins OLE1, 2, 4 and 5 (17, 18)(Figure S2A). Oleosins have been described to have a structural role as surfactants and to be seed and pollen-specific proteins (36). To our knowledge, they were so far not found in the proteome of vegetative tissues from flowering plants (33, 34). That said, *OLE* transcripts occur in yellow nutsedge tubers (11) and the leaves of the resurrection grass *Oropetium thomaeum*, after desiccation (9). Based on these findings and the presence of oleosin transcripts in *Physcomitrium* (*Physcomitrella*) *patens* spores and drought stressed algae, it is conceivable that oleosins are generally needed to shelter LDs under desiccation (5).

**Figure 3.**
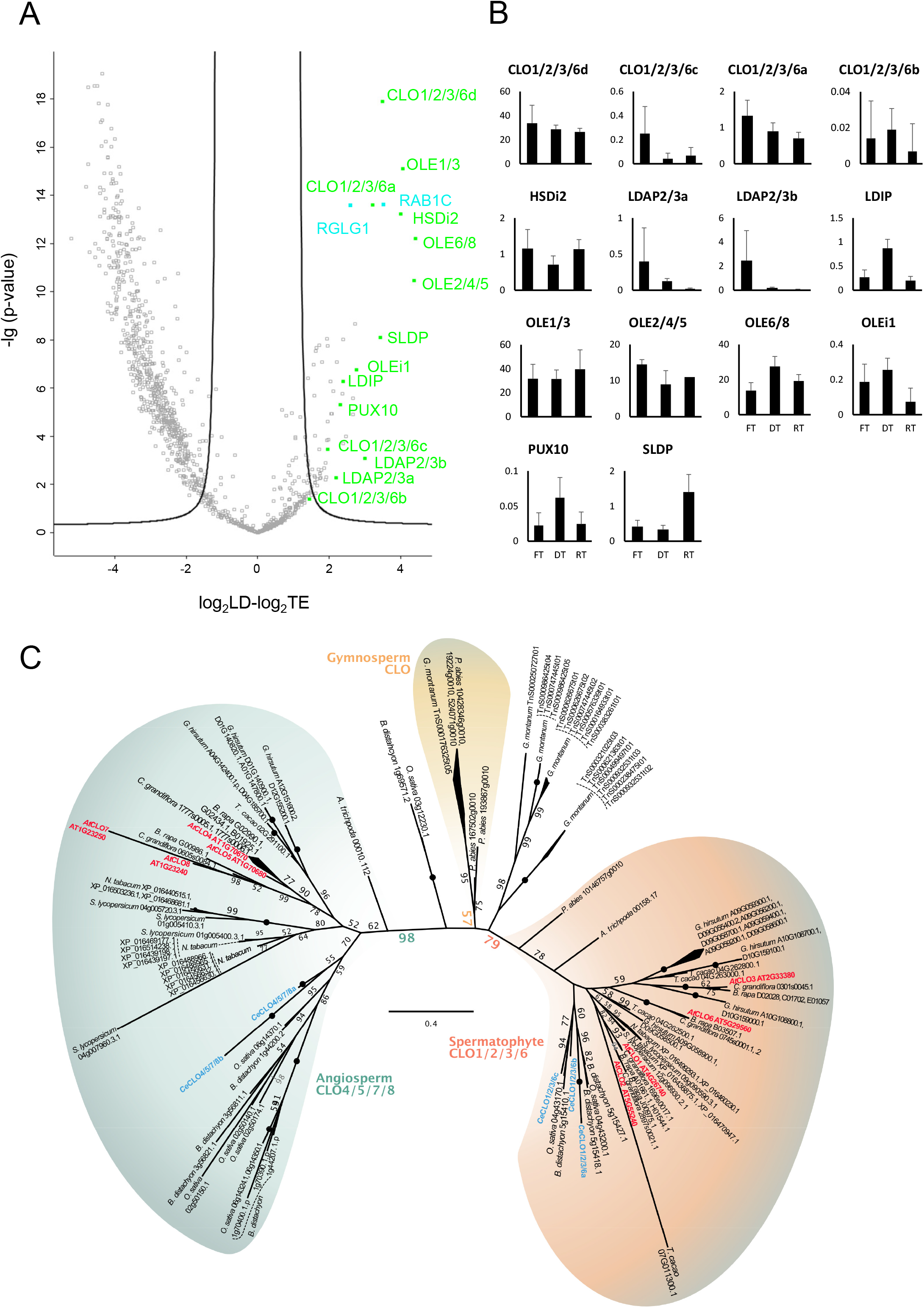
LD proteomes of yellow nutsedge tubers resemble those of seeds and seedlings. A volcano plot was constructed to visualize proteins, which are significantly and consistently LD-enriched in yellow nutsedge (A). All 15 LD-enriched samples were compared to all total cellular fractions of fresh, dry, and rehydrated tubers (Table S2). The log_2_-transformed values and p-values were calculated. Known LD proteins are indicated in green, while interesting candidates are in blue (see Table S8 for a list of all candidates). Black lines indicate a false discovery rate of 0.01. (B) The riBAQ intensities of LD-associated proteins in the LD-enriched fraction were calculated as a percentage of the riBAQ of all known LD-associated proteins (see Table S8 for data). (C) Unrooted maximum likelihood phylogeny of caleosin homologs detected in the predicted proteomes of spermatophyte genomes and the yellow nutsedge transcriptomes. Yellow nutsedge and Arabidopsis caleosins are depicted in blue and red, respectively. Abbreviations: CLO, caleosin; HSD, steroleosin; LDAP, LD-associated protein; LDIP, LDAP interacting protein; OLE, oleosin; PUX, plant UBX domain-containing protein; RAB, Rab GTPase; RGLG, RING domain ligase, SLDP, seed LD protein.

Steroleosins are likewise predominantly found in seeds and not in vegetative tissues (37). They are speculated to play a role in the metabolism of brassinosteroids (38, 39), hormones involved in development, and seed dormancy (40). There were two steroleosins isoforms in tubers (Figure S2B). This warrants future work on their role in development, dormancy or sprouting. Interestingly, seed lipid droplet protein (SLDP), another LD-associated protein (18) that is seed and seedling specific in Arabidopsis (17), was also detected in tubers but its molecular function is so far unknown.

Caleosins (CLO), in comparison, have been found in seeds, pollen and leaves (41-43). In Arabidopsis, CLO1 and CLO2 are the dominant caleosins of seeds (18); CLO3 is likely the most abundant leaf LD protein (34, 35) and is upregulated under stress (24). We explored whether the caleosins of yellow nutsedge showed a specific phylogenetic relationship to the seed or leaf isoforms of Arabidopsis. We however found that CLO1, 2 3 belongs to a larger clade within CLO gene family (here coined ‘CLO1/2/3/6’) within which lineage-specific duplications occurred; the most recent common ancestor of monocots and dicots likely had a single CLO1/2/3/6 homolog (Figure 3C).

Two further ubiquitously present LD proteins involved in the proper formation of LDs are the lipid droplet associated protein (LDAP) (44) and its interaction partner, LDAP interacting protein (LDIP) (45). LDAPs are low abundant in both desiccated tubers (Figure 3B) and seeds (18) while their interaction partner LDIP is also abundant in the desiccated structures of both species.

Apart from known LD proteins, also further proteins were found enriched in the LD fraction. Especially strongly enriched was a small G protein, RAB1C. Its mammalian homolog RAB18 is known to localize to LDs in mammals (46) but LD-binding G proteins are so far not described in plants. It appears possible, though that they are involved in LD formation or trafficking.

Another LD-enriched protein, the E3 ubiquitin ligase RGLG1 (RING domain ligase 1) (47), would be interesting to study in the future considering that several LD proteins get ubiquitinated prior to degradation (32, 48-50). RGLG1 from Arabidopsis is predominantly expressed in seeds (17) and was enriched in the LD-fractions of seedlings (18). It also plays a role in drought responses (47).

### Conclusions and outlook

There are striking similarities between Arabidopsis seeds and tubers of yellow nutsedge. It is therefore conceivable that oil accumulation and desiccation resistance of both organs is based on a similar proteome and ultimately a similar gene expression pattern that could be regulated by a similar network of hormones and transcription factors. This network simply shifted its tissue-specificity, resulting in the co-option of an established molecular program by a different organ. Indeed, comparable protein patterns might not have occurred first in seeds, but across land plants to sustain a variety of oil-rich desiccation tolerant structures (5). For example, also the oil rich and drought resistant spores of *P. patens* abound in transcripts of typical LD and LEA proteins (51, 52).

Apparently, the reprogramming of yellow nutsedge tubers to seed-like characteristics evolved during a comparable short time frame. This underlines the potential for a complete reprogramming of vegetative tissues in crop plants by genetic engineering (53-55). Strategies observed in tubers of yellow nutsedge have the potential to guide such biotechnological approaches.

## Material and methods

### Plant Material

Yellow nutsedge (*Cyperus esculentus* L. var. *sativus*) tubers were either taken dry, rehydrated for two days in water under continuous airflow or allowed to sprout in soil and harvested 1-2 days after sprouting. Yellow nutsedge plants were harvested from plants grown in a greenhouse under 14-16 h artificial light of mercury-vapor lamps (150 μmol m^−2^ sec^−1^) complemented with sunlight and a temperature of 20-22°C at daytime or 16-18°C at nighttime. Roots and leaves were harvested from 33 d old plants and fresh tubers from 4 month old plants.

### Isolation of total and LD-enriched protein fractions

For each condition, 5 biological replicates were processed. Biological replicates used for lipid droplet isolation comprises 5 individual tubers, tissues of germinated tubers, leaves and roots derived from single individuals. The tubers were ground in a precooled mortar with 1-2 g sea-sand and 10 ml grinding buffer (50 mM Tris pH 7.4, 10 mM KCl, 200 mM Sucrose, 200 μM PMSF). Subsequently, 2 ml of the suspension were transferred into a 2 ml reaction tube and spun for 10 s at 1000 g. A 50 μl aliquot, referred to as total extract, was taken from the supernatant into 1 ml ethanol. The remaining material was centrifuged for 20 min at 20,000 g at 4°C. The floating fat pad was mechanically picked with a spatula and washed three times in 1.7 ml fresh grinding buffer by 20 min at 20,000 g at 4°C centrifugation. The resulting crude fat pad was resuspended in 1 ml ethanol and referred to as LD-enriched fraction. Total extract and LD-enriched fraction were stored at −20°C for at least one day to enhance protein precipitation.

### Peptide sample preparation

The protein pellets were dissolved in 6 M urea and 5 % (w/v) SDS. Protein concentrations were determined with a Pierce BCA protein assay kit (Thermo Fisher Scientific, Waltham, MA, USA). 10 μg of protein was run on a SDS-PAGE gel until they entered the separation gel. A single gel piece per sample containing all proteins was excised, tryptic digested and derivatised (56). Peptides were desalted over an Empore™ Octadecyl C18 47 mm extraction disks 2215 (Supelco, St. Paul, MN, USA) as described (57).

### LC-MS/MS analysis

Dried peptide samples were reconstituted in 20 μl LC-MS sample buffer (2% acetonitrile, 0.1% formic acid). 3 μl of each sample were subjected to reverse phase liquid chromatography for peptide separation using an RSLCnano Ultimate 3000 system (Thermo Fisher Scientific). Therefore, peptides were loaded on an Acclaim PepMap 100 pre-column (100 μm × 2 cm, C18, 5 μm, 100 Å; Thermo Fisher Scientific) with 0.07% trifluoroacetic acid at a flow rate of 20 μL/min for 3 min. Analytical separation of peptides was done on an Acclaim PepMap RSLC column (75 μm × 50 cm, C18, 2 μm, 100 Å; Thermo Fisher Scientific) at a flow rate of 300 nL/min. The solvent composition was gradually changed within 94 min from 96 % solvent A (0.1 % formic acid) and 4 % solvent B (80 % acetonitrile, 0.1 % formic acid) to 10 % solvent B within 2 minutes, to 30 % solvent B within the next 58 min, to 45% solvent B within the following 22 min, and to 90 % solvent B within the last 12 min of the gradient. All solvents and acids had Optima grade for LC-MS (Thermo Fisher Scientific). Eluting peptides were on-line ionized by nano-electrospray (nESI) using the Nanospray Flex Ion Source (Thermo Fisher Scientific) at 1.5 kV (liquid junction) and transferred into a Q Exactive HF mass spectrometer (Thermo Fisher Scientific). Full scans in a mass range of 300 to 1650 m/z were recorded at a resolution of 30,000 followed by data-dependent top 10 HCD fragmentation at a resolution of 15,000 (dynamic exclusion enabled). LC-MS method programming and data acquisition was performed with the XCalibur 4.0 software (Thermo Fisher Scientific).

### Generation of protein library of *Cyperus esculentus* for peptide identification

Total RNA was extracted from frozen tissues of *Cyperus esculentus* mixed tissue sample and 8 different stages of tuber development by homogenizing material in Plant RNA Reagent according to instructions (Invitrogen, Carlsbad, USA). RNA integrity and concentration were determined using Experion RNA StdSens analysis kit (BioRad, Hercules, USA). Total RNA was DNase treated (TurboDNase, Ambion, Carlsbad, USA) before sequencing. Libraries for RNAseq were prepared at BGI (Shenzhen, China). Libraries were sequenced through the Illumina sequencing platform HiSeq 2000 as unpaired-end reads. Two different software were used to predict open-reading frames or proteins potentially encoded by the produced contigs (putative transcripts). TransDecoder (https://github.com/TransDecoder) Release v5.0.1 (58) was applied using default settings. GeneMark.hmm eukaryotic (http://exon.gatech.edu/GeneMark/gmhmme.cgi) was applied (59) using Arabidopsis as species model and with protein sequence as selected output for processing of the contigs. An additional protein reference was based on a previously published yellow nutsedge transcriptome (11). ORFs were extracted and translated using Geneious 8.1.8 (Biomatters Ltd., Auckland, New Zealand).

### Data processing Calculation of relative intensity-based absolute quantification (riBAQ) values

MS data was processed with Max Quant software version 1.6.2.10 (60, 61) with standard settings except for: „Match between runs‟ was turned on; „iBAQ‟ was selected for label free quantification; FTMS recalibration was turned on. The three above mentioned nutsedge protein libraries were used. iBAQ values were determined with Max Quant 1.6.2.10. The data was further processed with Perseus software version 1.6.2.2 (62). Reverse hits, contaminants and proteins only identified by a modified peptide were removed from the data matrix. Then, all iBAQ values were divided by the total iBAQ value per sample and multiplied by 1000 yielding riBAQ in per mill. These values were the basis for all further data analysis.

### BLAST

Homologs of nutsedge proteins in *Arabidopsis thaliana* were detected by using a BLASTp approach (63, 64). For this, protein data predicted from nutsedge transcriptomes were used as queries against the *Arabidopsis thaliana* TAIR10 primary transcript protein release using BLAST 2.5.0+.

### PCA

The PCA of the nutsedge total extracts and the comparison of total and LD enriched fractions was performed on riBAQ values. Only proteins were considered that were identified by a least two peptides and that were found in at least three replicates of one of the stages. For the PCA comparing yellow nutsedge and Arabidopsis data, only proteins were considered that had a homolog in both species and were found in four out of five replicates of at least one stage. The values for each protein was normalized within the individual species setting the average of the highest stage to 1. PCA plots were created with Perseus 1.6.2.2 (62) using standard settings (Category enrichment in components was turned off).

### Hierarchical clustering

The clustering was performed on average values for each stage/tissue. Furthermore, the highest average of each protein within each species was set to 1. Only proteins were considered that were found in all samples of at least one stage. In the comparison of yellow nutsedge and Arabidopsis only proteins were incorporated that had a homolog in both species. Plots were created with Perseus 1.6.2.2 (62) using Euclidian distance, average linkage and no constrains. The data was preprocessed with k-means. The number of clusters was set to 300, the maximum number of iterations to 100 and the number of restarts to 100. In the end, the proteins were grouped in 30 clusters.

### Pearson correlation

Pearson correlation was performed on average values for each stage/tissue. Furthermore, the highest average of each protein within each species was set to 1. Only proteins were considered that were found in all samples of at least one stage and that had a homolog in both species. The correlation coefficients were calculated with Perseus 1.6.2.2 (62).

### GO term enrichment

Go term enrichment analysis of proteins contained in certain clusters was done using an online tool based on the PANTHER classification system (http://geneontology.org/; http://pantherdb.org/webservices/go/overrep.jsp) (16).

### Quantitative GO-term analysis

The quantitative GO-term analysis was based on riBAQ values. First, yellow nutsedge Proteins were assigned Arabidopsis identifiers if the e-value of the BLASTp result was lower than 10^−5^. If several yellow nutsedge proteins were assigned the same Arabidopsis identifier, the values were added. Then, all proteins were assigned one or more GO terms and the values added for each term.

### Volcano plots

Only proteins were considered that were found in four out of five replicates of at least one stage/subcellular fraction. Missing values were imputed with Perseus 1.6.2.2 (62) using a width of 0.3 and a downshift of 1.8 separately for each column. The volcano plot was then created using a t-test, a number of 250 randomizations, a false discovery rate of 0.01 and an S_0_ of 2.

### Analysis of LD proteins

All proteins that were homolog to a known LD protein from Arabidopsis were considered as LD proteins. The abundance of each LDs protein was divided by the total abundance of all LD proteins in the sample.

### Phylogenetic analyses

In order to construct phylogenies, the datasets of the proteins of interest were supplemented with additional homologs from other seed plants and, in case of LDAPs, land plants and green algae. For this, well-described *Arabidopsis thaliana* proteins were used as query sequences in a BLASTp search against protein data from genome releases. In each case, the detected protein homologs in the yellow nutsedge transcriptomes were added.

For the phylogenetic analysis of CLO and HSD, homologs were mined from genome data of *Theobroma cacao* (65), *Picea abies* (66)*, Oryza sativa* (67)*, Nicotiana tabacum* (68),*Gnetum montanum* (69)*, Gossypium hirsutum* (70)*, Capsella grandiflora* (71)*, Brassica rapa* (72)*, Brachypodium distachyon* (73)*, Amborella trichopoda* (74)*, Arabidopsis thaliana* (15), and *Solanum lycopersium* (75).

For the phylogenetic analysis of OLE, homologs were mined from genome data of *Oryza sativa* (67)*, Arabidopsis thaliana* (15), and *Solanum lycopersium* (75).

For the phylogenetic analysis of LDAPs, the same selection of homologs as in de Vries and Ischebeck (2020) were chosen, i.e. *Theobroma cacao* (65)*, Triticum aestivum* (76)*, Selaginella moellendorffii* (77)*, Solanum lycopersium* (75)*, Physcomitrium patens* (78)*, Picea abies* (66)*, Oryza sativa* (67)*, Nicotiana tabacum* (68)*, Marchantia polymorpha* (79)*, Gnetum montanum* (69)*, Gossypium hirsutum* (70)*, Carica papaya* (80)*, Capsella grandiflora* (71)*, Brassica rapa* (72)*, Brassica oleracea* (81)*, Brachypodium distachyon* (73)*, Amborella trichopoda* (74)*, Arabidopsis thaliana* (15)*, Arabidopsis lyrata* (82), the hornworts *Anthoceros agrestis* and *Anthoceros punctatus* (83), the ferns *Azolla filiculoides* and *Salvinia cucullata* (84), the Charophyceae *Chara braunii* (85), the genomes of the Zygnematophyceae *Mesotaenium endlicherianum* and *Spirogloea muscicola* (86) and the transcriptomes of the filamentous Zygnematophyceae *Zygnema circumcarinatum* SAG2419 (87) and SAG698-1a (88) as well as *Spirogyra pratensis* MZCH10213 and *Mougeotia* sp. MZCH240 (89), and genomes of the early-diverging streptophyte algae *Mesostigma viride* and *Chlorokybus atmophyticus* (90)

All protein sequences were aligned using MAFFT v7.453 L-INS-I (91). Sequences were cropped to retain the conserved region/protein features and re-aligned; sequences that did not cover at least 50% of the conserved region were removed. Maximum likelihood phylogenies were computed using IQ-TREE (92) multicore version 1.5.5 for Linux 64-bit. 500 bootstrap replicates were computed. Each run of IQ-TREE entailed determining the best model for protein evolution according to the Bayesian information criterion (BIC) score via ModelFinder (93). The best models were: LG+G4 for CLO, JTT+G4 for OLE and HSD, and JTT+|+G4 for LDAP.

## Acknowledgements

PWN, OV, KS, GB and TI thanks the German research foundation (DFG, Grants IS 273/7-1, IRTG 2172 PRoTECT, BR1502-15-1 and INST 186/1230-1 FUGG to Stefanie Pöggeler) and JdV the European Research Council for funding (Grant Agreement No. 852725; ERC-StG ‘TerreStriAL’ under the European Union’s Horizon 2020 research and innovation programme). OV and KS on behalf of the Service Unit LCMS Protein Analytics of the Göttingen Center for Molecular Biosciences (GZMB) thank the DFG for funding (INST 186/1230-1 FUGG to Stefanie Pöggeler). PH was financed by grants from the Swedish Foundation for Strategic Research (SSF) and Trees and Crops for the Future (TC4F), a Strategic Research Area at SLU, supported by the Swedish Government.

## Author contributions

P.W.N., G.H.B., P.H. and T.I. designed the work, P.W.N., K.S., O.V., J.d.V., A.S.C., P.H. and T.I. performed research, P.W.N., K.S., G.H.B., J.d.V., A.S.C., P.H. and T.I. analyzed data, and P.W.N., K.S., J.d.V., P.H., and T.I. wrote the manuscript. All authors critically read and revised the manuscript and approved the final version.

## Supplemental information

**Figure S1.** Yellow nutsedge developmental stages investigated in this study.

**Figure S2.** LD enriched fractions could be reproducibly isolated.

**Figure S3.** Oleosin (OLE) and steroleosin (HSD) phylogeny.

**Figure S4.** Phylogeny of lipid droplet associated proteins (LDAPs).

---

**Figure S1.**
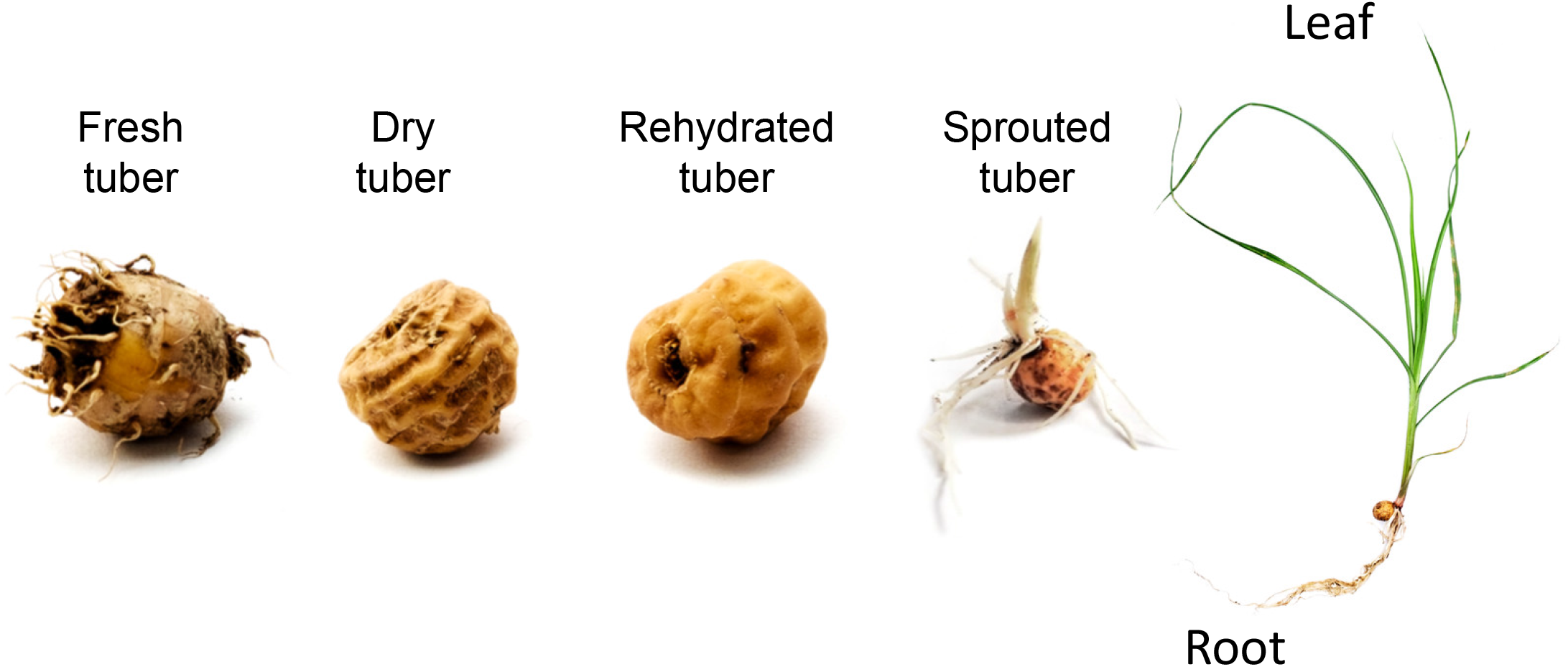
Yellow nutsedge developmental stages investigated in this study. Rehydrated tubers were soaked in water for 48 h and tubers sprouted in the soil were harvested 1–2 days after onset of sprouting. Roots and leaves were taken from plants grown in soil for 33 days omitting the tuber.

**Figure S2.**
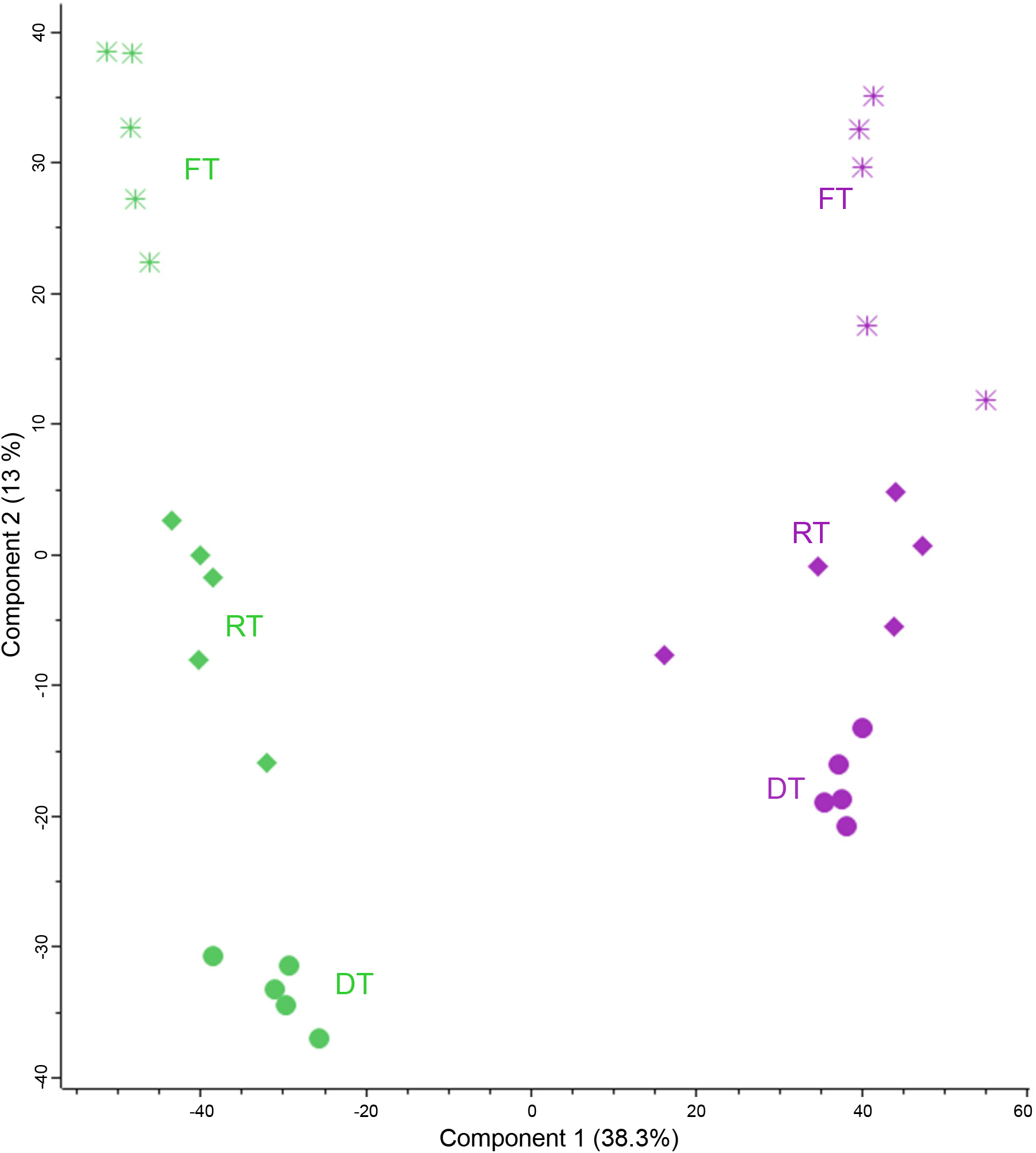
LD enriched fractions could be reproducibly isolated. Total protein extracts and LD-enriched fractions of yellow nutsedge tubers (fresh, FT; dry, DT, rehydrated for 48 h (RT) were compared in respect to their proteome (Table S2). A principal component analysis was performed with all proteins that were found in at least three replicates of one stage (A). n=5 biological replicates per stage and fraction.

**Figure S3.**
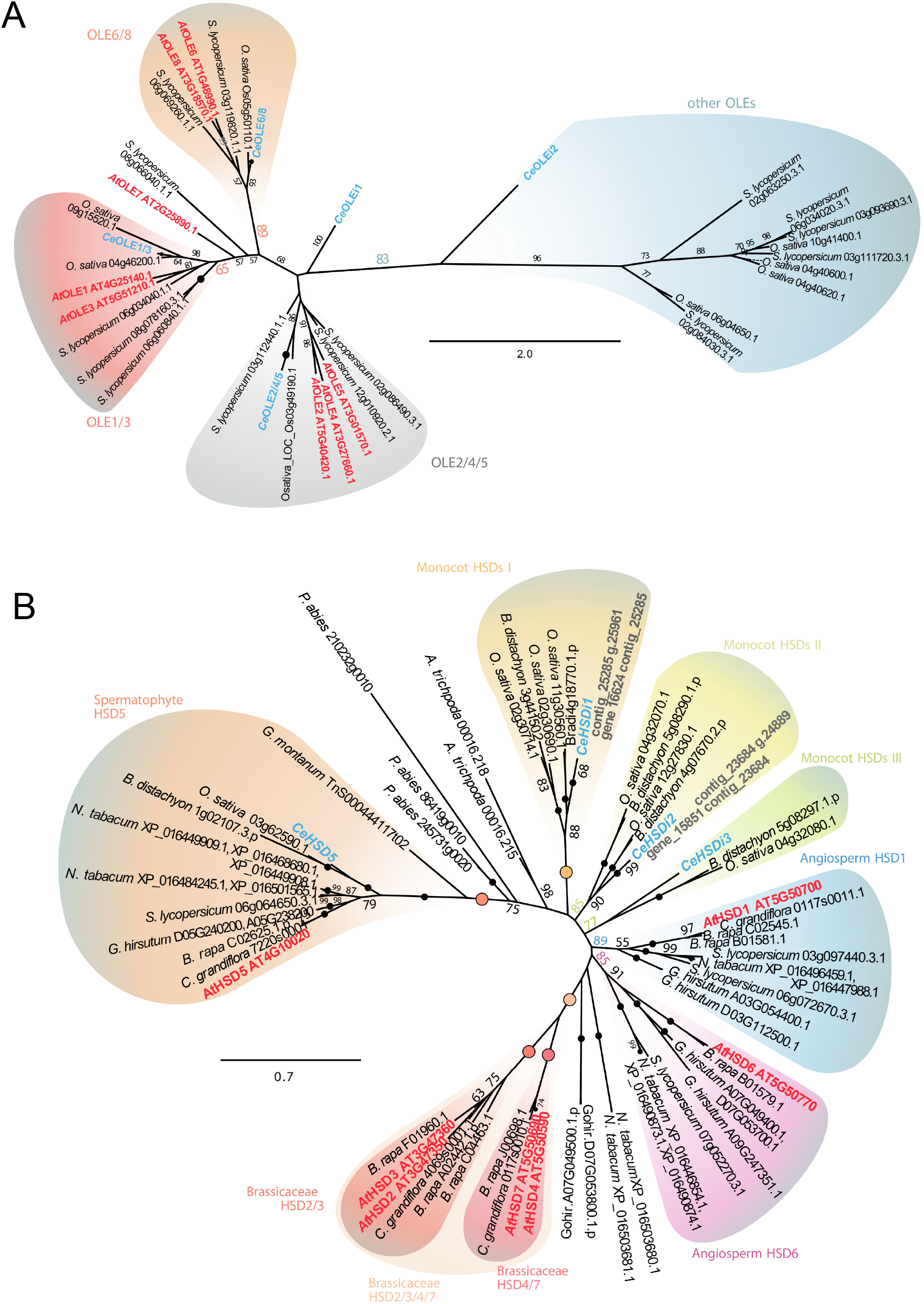
Oleosin (OLE) and steroleosin (HSD) phylogeny. Unrooted maximum likelihood phylogeny of oleosin (A) and steroleosin (B) homologs detected in the predicted proteomes of spermatophyte genomes and the yellow nutsedge transcriptomes. Yellow nutsedge and Arabidopsis proteins are depicted in blue and red, respectively.

**Figure S4.**
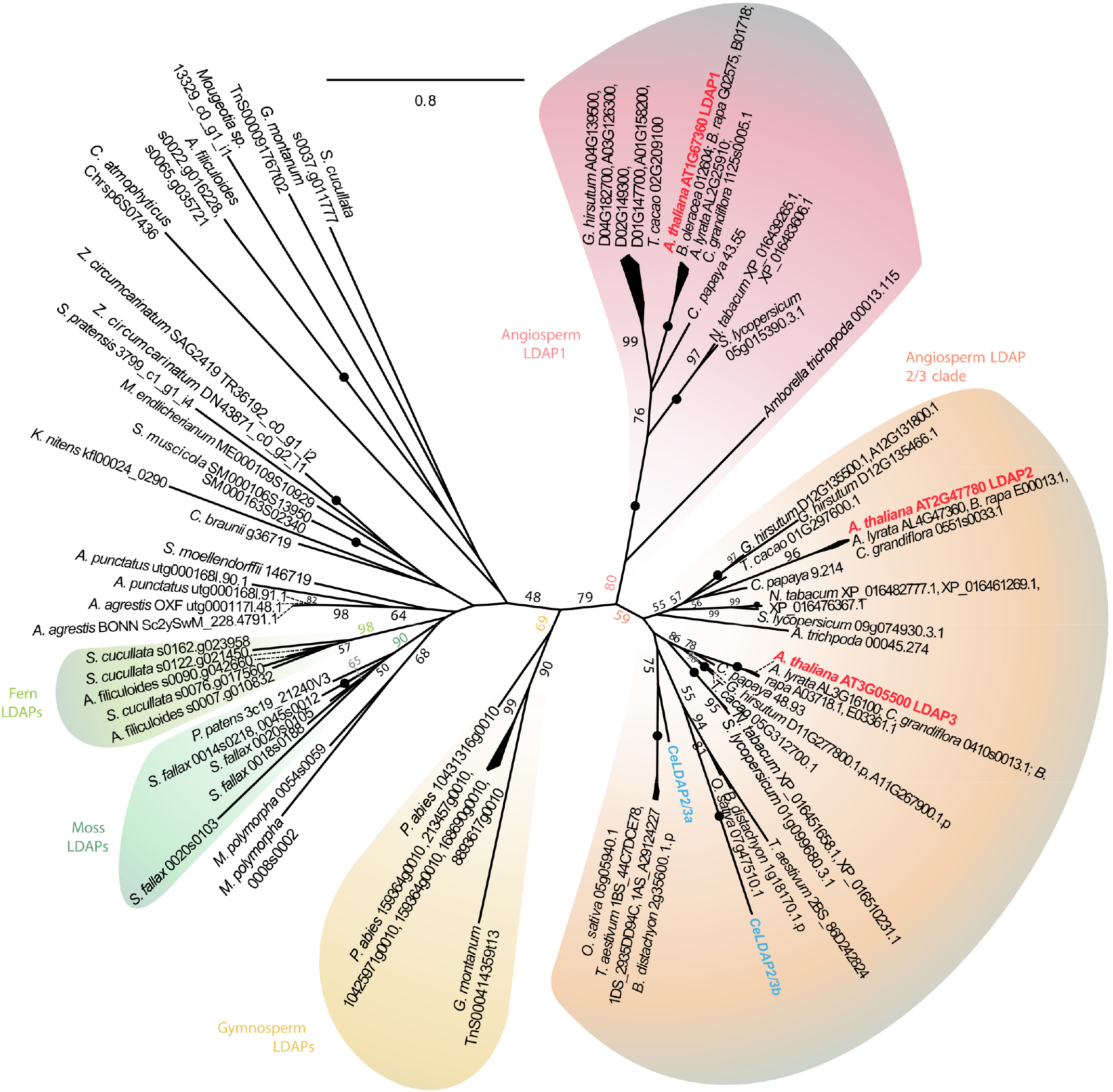
Phylogeny of lipid droplet associated proteins (LDAPs). Unrooted maximum likelihood phylogeny of homologs detected in the predicted proteomes of spermatophyte genomes and the yellow nutsedge transcriptomes. Yellow nutsedge and Arabidopsis proteins are depicted in blue and red, respectively.

